# A whole-genome CRISPR screen identifies the spindle accessory checkpoint as a locus of nab-paclitaxel resistance in pancreatic cancer cells

**DOI:** 10.1101/2024.02.15.580539

**Authors:** Priya Mondal, George Alyateem, Allison V. Mitchell, Michael M. Gottesman

## Abstract

Pancreatic adenocarcinoma is one of the most aggressive and lethal forms of cancer. Chemotherapy is the primary treatment for pancreatic cancer, but resistance to the drugs used remains a major challenge. A genome-wide CRISPR interference and knockout screen in the PANC-1 cell line with the drug nab-paclitaxel has identified a group of spindle assembly checkpoint (SAC) genes that enhance survival in nab-paclitaxel. Knockdown of these SAC genes (BUB1B, BUB3, and TTK) attenuates paclitaxel-induced cell death. Cells treated with the small molecule inhibitors BAY 1217389 or MPI 0479605, targeting the threonine tyrosine kinase (TTK), also enhance survival in paclitaxel. Overexpression of these SAC genes does not affect sensitivity to paclitaxel. These discoveries have helped to elucidate the mechanisms behind paclitaxel cytotoxicity. The outcomes of this investigation may pave the way for a deeper comprehension of the diverse responses of pancreatic cancer to therapies including paclitaxel. Additionally, they could facilitate the formulation of novel treatment approaches for pancreatic cancer.

## Introduction

Pancreatic cancer is characterized by its aggressive nature and resistance to conventional therapies^1^. A first-line treatment for pancreatic cancer is a combination of nab-paclitaxel and gemcitabine^2^. Paclitaxel stabilizes microtubules by binding tightly to the β-subunit of α/β-tubulin dimers, inhibiting their dynamic function^3^, but the precise mechanism of its cytotoxicity is not known^4^. Nab-paclitaxel (tradename Abraxane) is a formulation of paclitaxel with albumin. It is created by homogenizing 3-4% serum albumin with paclitaxel to improve drug biodistribution^5^. Paclitaxel is also used in combination with cisplatin or carboplatin for treating various types of primary cancers^6^. In recurrent cancer cases, a dose-intensive paclitaxel regimen alone is used as a second-line drug. However, the effectiveness of paclitaxel decreases with subsequent treatments, indicating the development of drug resistance^7^. Acquired paclitaxel resistance can occur by several mechanisms including overexpression of ABC transporters (especially ABCB1^8^), altered expression of apoptotic genes, changes associated with microtubules, as well as the break-up of cancer cell nuclei into micronuclei^7,9^. Therefore, understanding the mechanisms underlying paclitaxel resistance is crucial for the development of more effective therapies.

High-throughput genetic screening is a common method to investigate drug resistance in human cancers.

The development of the CRISPR/Cas9 system has significantly accelerated functional genomic research. In bacteria, this system acts as an RNA-based immune system, while in eukaryotic cells, it has been modified to create frameshift mutations and deletions for specific gene knockouts and to interfere with transcription by activating or inhibiting gene expression. A single guide RNA (sgRNA) guides Cas9 to the target sequence, causing a double-strand break in the target sequence. In this way, it is possible to delete or insert the desired sequence into genes. If the Cas9 nuclease is inactivated and specific transcriptional activators or inhibitors are fused to it, specifically targeted genes can be turned on or off. Various CRISPR/Cas9 libraries have recently been developed for genetic screening in mammalian cell culture and mouse models. These CRISPR/Cas9 library screens have been used to identify genes that play important roles in cancer cell survival, proliferation, migration, and resistance to drug treatment in various models^10-13^.

In this study, we perform a genome-wide CRISPR/Cas9 interference and knockout screen in pancreatic adenocarcinoma (PANC-1) cells with nab-paclitaxel to identify genes that promote survival of cells treated with nab-paclitaxel. We found three important spindle assembly checkpoint (SAC) regulators (BUB1B, BUB3, TTK), among the top hits of the screen. These proteins play a crucial role in ensuring accurate chromosome segregation during cell division.

It was observed that knockdown of these SAC genes reduced the cytotoxic effect of nab-paclitaxel. Small molecule inhibitors that target the phosphorylation of TTK and diminish its kinase activity were also found to antagonize the effect of nab-paclitaxel. These studies suggest that the SAC regulators are critical mediators of paclitaxel efficacy in PANC-1 cells.

## Methods

### Chemicals

Drugs were obtained as follows: Nab-paclitaxel (Abraxane, Bristol Myers Squibb), paclitaxel (Sigma T7191), vincristine sulfate (Sigma V8879), BAY 1217389 (Selleckchem.com S8215), MPI0479605 (Selleckchem.com S7488), BAY 1816032 (ChemieTek CT-BAY181).

### Cell culture

PANC-1 (ATCC) cells were cultured in DMEM (Life Technologies) with 10% fetal bovine serum (Life Technologies), 1% penicillin-streptomycin, and 1% glutamine (Life Technologies). Cells were incubated at 37°C, in 5% CO2, and passaged twice a week using Trypsin-EDTA (0.25%) (Life Technologies). Cells were tested for mycoplasma contamination using the MycoAlert Mycoplasma Detection Kit (Lonza) according to the manufacturer’s instructions.

### Genome-wide CRISPR/Cas9 interference and knockout library screen

The human CRISPR Brunello lentiviral pooled library (Addgene # 73178-LV)^14^ and human CRISPR Dolcetto (Set A) inhibition library (Addgene # 92386-LV)^15^ were used to identify genes responsible for enhanced survival of PANC-1 cells treated with nab-paclitaxel. The Brunello library contains 76,441 sgRNAs targeting 19,114 genes and 1000 non-targeted sgRNAs as a control. The Dolcetto (Set A) library contains 57,050 sgRNAs targeting 18,901 genes and 500 unique non-targeting controls. A schematic diagram of this genetic screen appears in our previous publication^13^. To obtain a cell-efficient knockout and interference library on the lentiviral vector, two stable PANC-1 cell lines were established by lentiviral transduction, one expressing Cas9 (Addgene # 52962-LV)^16^ and one expressing dCas9-KREB (Addgene # 89567)^17^. The expression of Cas9 and dCas9 were confirmed by Western blotting (Fig. S1A, B). PANC-1-Cas9 and PANC-1-dCas9-KREB cells were transduced with the Brunello and Dolcetto libraries^15^, respectively, at a low MOI (∼0.3) to ensure effective barcoding of individual cells. The transduced cells were selected with 2⍰μg/ml of puromycin for 3 days to generate a cell pool carrying the libraries, which was then treated with vehicle (DMSO) and nab-paclitaxel (10⍰µM) for 10 days, respectively. After treatment, at least 1⍰×⍰10^9^ cells were collected to ensure over 500 × coverage of the library. Genomic DNA was extracted using the QIAmp DNA blood Cell Maxi Kit (Qiagen) according to the manufacturer’s protocol. The sgRNA sequences were amplified using Taq polymerase (Takara Bio, Inc.) and adapted for sequencing. The desired DNA product was purified with 6% TBE gel (Invitrogen) and subjected to massive parallel amplicon sequencing carried out by an Illumina sequencer (Sanger/Illumina 1.9). The sgRNA read count and hits calling were analyzed by the MAGeCK v0.5.7 algorithm^18^.

### Lentiviral transduction

Knockdown of the BUB1B, BUB3 and TTK genes in PANC-1 was achieved by transfecting cells with short hairpin RNA (shRNA)-packed lentiviral particles (Santa Cruz Biotechnology). In a 6-well plate, 1 × 10^5^ cells were transduced (MOI 1) with shRNA-packed lentiviral particles against human BUB1B (sc-37542-V), BUB3 (sc-37540-V), TTK (sc-36758-V) or a non-targeting control (sc-108080) in triplicate using Polybrene Transduction Reagent (5 µg/ml; Millipore). Single-cell colonies were subsequently isolated by selection with puromycin (5 µg/ml; InvivoGen) and the phenotype was analyzed by Western blot.

### Plasmid transfection

Human BUB1B, BUB3, and TTK genes were introduced in PANC-1 using pcDNA3.1+/C-(K)-DYK (GenEZ; GenScript) vectors (10 µg/ml). Lipofectamine was used as a transfection reagent (Life Technologies). OPTIMEM (Invitrogen) was used as transfection media. Single clones were isolated after selection with G418 (2mg/ml; Corning). Overexpression phenotype was confirmed by Western blot.

### Cytotoxicity assays

Cells were seeded in opaque white flat bottom 96-well plates at 1000 cells/well density and allowed to attach overnight. Cells were then treated with increasing concentrations of the desired compound in complete media and incubated for the desired time. The cell growth inhibition was determined using Cell Titer Glo (Promega) according to the manufacturer’s instructions. Luminescence was recorded in a microplate reader (Tecan Infinite M200 Pro) for 100 ms integration time.

### Cell growth rate analysis

Kinetic cell proliferation assays were monitored using the IncuCyte S3 Live Cell Analysis System (Essen Bioscience). 12-well plates were incubated at 37°C, in 5% CO2. Sixteen non-overlapping planes of view phase-contrast images were captured using a 10x objective, with data collected every 4 hr for the duration of each experiment. Incucyte Base Software was used to calculate average confluence. Population doublings were calculated using the formula Tdoubling = (log2(ΔT))/(log(c2) ™ log(c1)), where c1 and c2 are the minimum and maximum percentage confluency during the linear growth phase, respectively, and ΔT was the time elapsed between c1 and c2.

### Western blotting

Cells were harvested post-treatment and re-suspended in lysis buffer (50⍰mM Tris-HCl pH 7.4, 150⍰mM NaCl, 1% NP-40, 1% protease inhibitor cocktail (Cell Signaling), sonicated, and centrifuged to remove cell debris. The supernatant (30 µg protein) was separated by 4–12% Bis-Tris NuPAGE gel (ThermoFisher) and protein size was estimated using the Precision Plus All Blue (BioRad) ladder. Separated protein was transferred to a 0.2⍰μm pore nitrocellulose membrane (VitaScientific) using a wet transfer electrophoresis system (Life Technologies). The resulting membrane was blocked in Odyssey PBS Blocking Buffer (LI-COR) for 1⍰h at room temperature and subsequently incubated, with the respective primary antibodies diluted in blocking solution overnight at 4°C with gentle agitation. Membranes were then washed with 0.5X TBS (KD Medical) containing 0.5% TBS-Tween-20 (Boston BioProducts) three times for 15 min prior to and after the addition of an IRDye Goat anti-Rabbit (LI-COR) or anti-Mouse (LI-COR) secondary antibody. Proteins were visualized using an Odyssey CLx imaging system (LI-COR). Relative expression of proteins were quantified using Image Studio Lite Quantification Software (LI-COR). The following primary antibodies were used for Western blotting; BUB1B (1:1000, Cell Signaling; 4116S), BUB3 (1:1000, Cell Signaling; D8G6), TTK (1:1000, Cell Signaling; 3255S) and anti-GAPDH (1:8000; American Research Products, Inc; 6C5).

### Cell cycle assay

2.5 × 10^5^ cells were seeded in 6 well plates. To synchronize the growth, 2.5 mM thymidine (Sigma Aldrich) was incubated with the cells for 16 h and 22 h, respectively, at 8 h intervals. Cells were collected post-treatment from 6-well plates by trypsinization. Cell pellets were stained with a solution containing RNaseA (200 U/ml; Invitrogen), propidium iodide (0.1 mg/ml, Sigma Aldrich), and 0.1% Triton X (Sigma-Aldrich) for 30 min in the dark at room temperature. The cell cycle distribution was detected by a FACS Canto II flow cytometer (BD Biosciences) and data were analyzed with ModFit LT software.

### Apoptosis assay

All cell lines were plated at a density of 2.5 × 10^5^ cells per well in a 6-well plate and treated with BAY 1217389 and nab-paclitaxel for 48⍰h. Cell pellets were stained with FITC Annexin V (Bio Legend) and propidium iodide (Invitrogen) for 30 min. Samples were read using a FACS Canto II Flow Cytometer and FlowJo software was used to determine the percentage of annexin/PI positive cells.

### Microscopy

Cells were grown on plain glass coverslips (Zeiss). For analysis, cells were fixed in 4% paraformaldehyde (Electron Microscopy Science) and permeabilized with PBS (Gibco) containing 0.5% Triton X-100 (Sigma Aldrich). Cells were blocked with 4% blocking buffer (Cell Signaling) and then incubated with the specified primary antibodies. The following primary antibodies were used for immunofluorescence: anti-alpha-tubulin, clone DM1A, Alexa Fluor 555 (Conjugate) (1:100; Millipore 05-829-AF555), and Alexa Fluor 647 rat anti-Histone H3 (pS28) (1:100; BD 558217). Coverslips were mounted onto slides using Prolong Gold anti-fade mounting medium with DAPI (Invitrogen). Images were acquired with a microscope (Zeiss LSM880 Airyscan microscope) using ZEN 2.3 SP1 software. Images of z stacks with 0.3 - 0.9 μm steps covering the entire volume of the mitotic apparatus were collected with a Plan-Apochromatic 63x/1.4 Oil DIC M27.

### Statistical analysis

All data are expressed as mean⍰±⍰standard error of the mean of 3 to 6 experiments. Statistical differences among the groups were analyzed by one-way ANOVA and the Student’s t-test. Data were considered significant at P⍰< ⍰0.05.

## Results

### Whole genome CRISPR knockout and interference screens indicate that reduced expression of three spindle checkpoint regulators (BUB1B, BUB3, TTK) enables PANC-1 cell survival in nab-paclitaxel

To identify key genes involved in nab-paclitaxel resistance in pancreatic adenocarcinoma, we conducted high-throughput CRISPR/Cas9 pooled cell fitness screens in PANC-1 pancreatic adenocarcinoma cells.

We used the human CRISPR Dolcetto (Set A) library or the human CRISPR Brunello library for the interference or knockout screens, respectively. Our goal was to identify genes in which knockout or suppression resulted in an increased resistance to nab-paclitaxel through positive selection. Fitness screens were carried out with either vehicle (DMSO) or nab-paclitaxel for 10 days (Fig. 1A, B). Our findings showed that nab-paclitaxel treatment at 10 μM significantly suppressed cell proliferation when compared to the control group, indicating significant selection pressure (Fig. S1C). Using this CRISPR/Cas9 knockout and interference library screening, we identified a subset of sgRNAs targeting 985 and 993 genes that were significantly enriched (P < 0.05) in nab-paclitaxel-treated cells when compared to the control (Tables S1-S4). We found BUB1B and TTK were the most significant positively selected hits following nab-paclitaxel treatment in the CRISPR interference screen, while BUB3 was the top hit in the knockout screen. The administration of nab-paclitaxel, coupled with targeted knockdowns of the genes BUB1B, TTK, and BUB3 using sgRNA, led to a notable enhancement in cell survival, suggesting that the loss of these genes in PANC-1 cells promoted cell survival after nab-paclitaxel treatment under our selection conditions (Fig. 1C, D).

**Figure 1.**
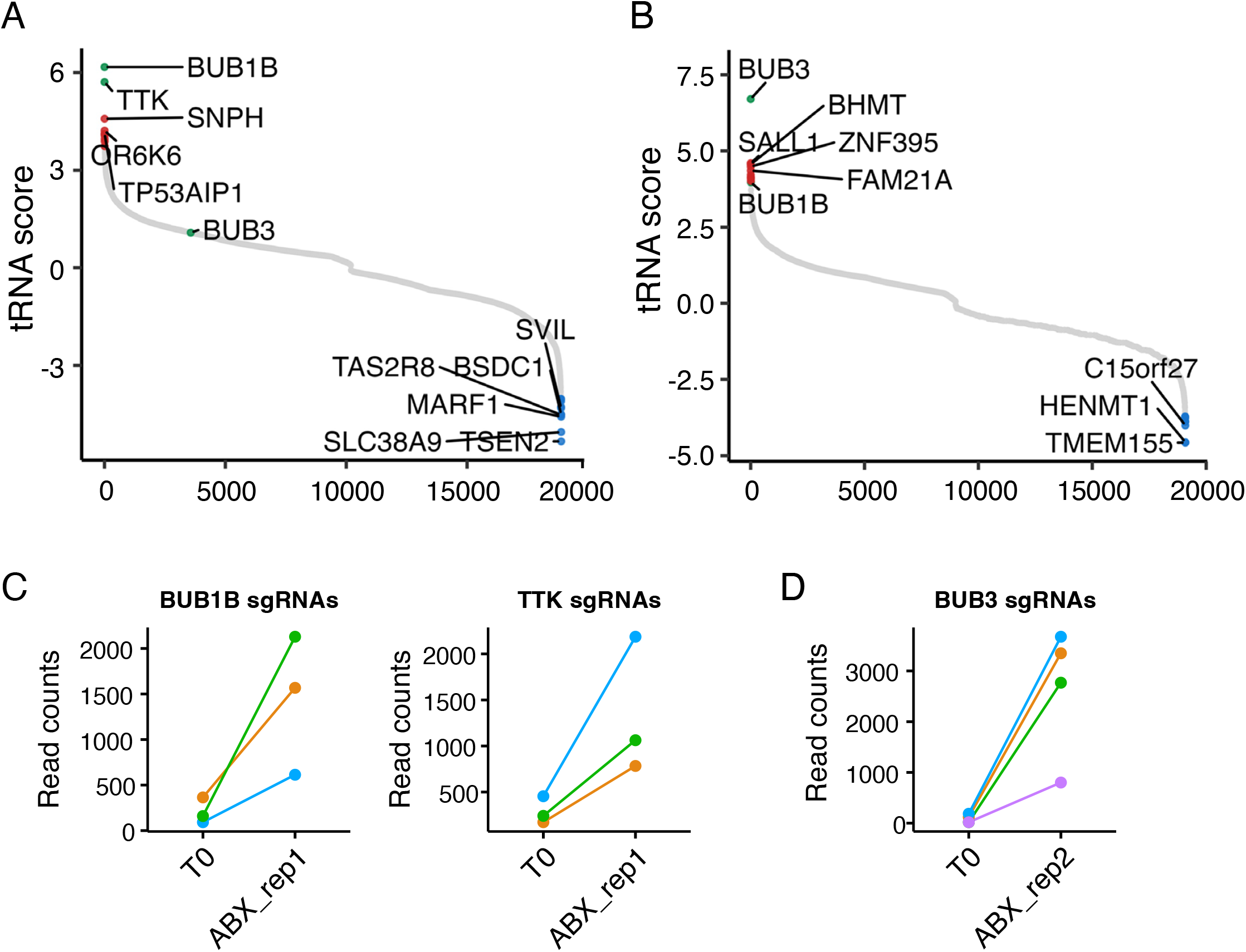
Genome-wide CRISPR interference and knockout screens identify BUB1B, TTK, and BUB3 as drivers for nab-paclitaxel resistance in PANC-1 cells. (A, B) The Rank plot of genes generated by differential RRA score identifies BUB1B and TTK as the most significant genes in the CRISPR knockdown screen and BUB3 as the most significant gene in the CRISPR knockout screen. (C, D) The sgRNAs targeting BUB1B, TTK, and BUB3 were consistently enriched in nab-paclitaxel-treated cells. Source data are provided as a Source Data file.

### Inhibition of selective SAC genes decreases the sensitivity of PANC-1 cells to Abraxene and other microtubule-targeting drugs

To validate the results from the CRISPR screens, we created individual BUB1B, BUB3, and TTK stable shRNA knockdown (KD) subclones and matched nontargeted (NT) shRNAs in PANC-1 cells. We confirmed sufficient knockdown efficiency for each gene by Western blot (Fig. S2; each panel shows 1 clone out of 9 that were obtained). The effects of gene knockdown on response to nab-paclitaxel treatment were measured by the Cell Titer-Glo luminescent cell viability assay (Promega). We observed that knockdown of BUB1B, BUB3B, or TTK significantly increased the resistance of PANC-1 cells to nab-paclitaxel with an approximately 10-fold increase in the IC_50_ values (∼1 nM vs. 10 nM) (Fig. 2A). Similar effects were observed when cells were treated with paclitaxel; however, BUB3 KD showed a slightly diminished resistant phenotype compared to BUB1B and TTK knockdowns (Fig. 2B). Docetaxel (brand name Taxotere) is also an effective first-line therapy for breast cancer, as it stabilizes microtubules^19^. In this study, BUB1BKD, BUB3KD, and TTKKD PANC-1 clones also showed significant resistance to docetaxel compared to control shRNA-transfected cells (Fig. 2C). Vincristine (a *Vinca* alkaloid) prevents cell growth by attaching to tubulin and disrupting microtubule polymerization. It is commonly used in combination with other drugs for cancer treatment^20^. The selected SAC knockout PANC-1 clones also showed significant resistance to vincristine (Fig. 2D), suggesting that the basis of resistance is not the stabilization of microtubules, but the downstream effect of microtubule dysfunction caused by both paclitaxel, a microtubule-stabilizing agent, and vincristine, a microtubule-destabilizing agent.

**Figure 2.**
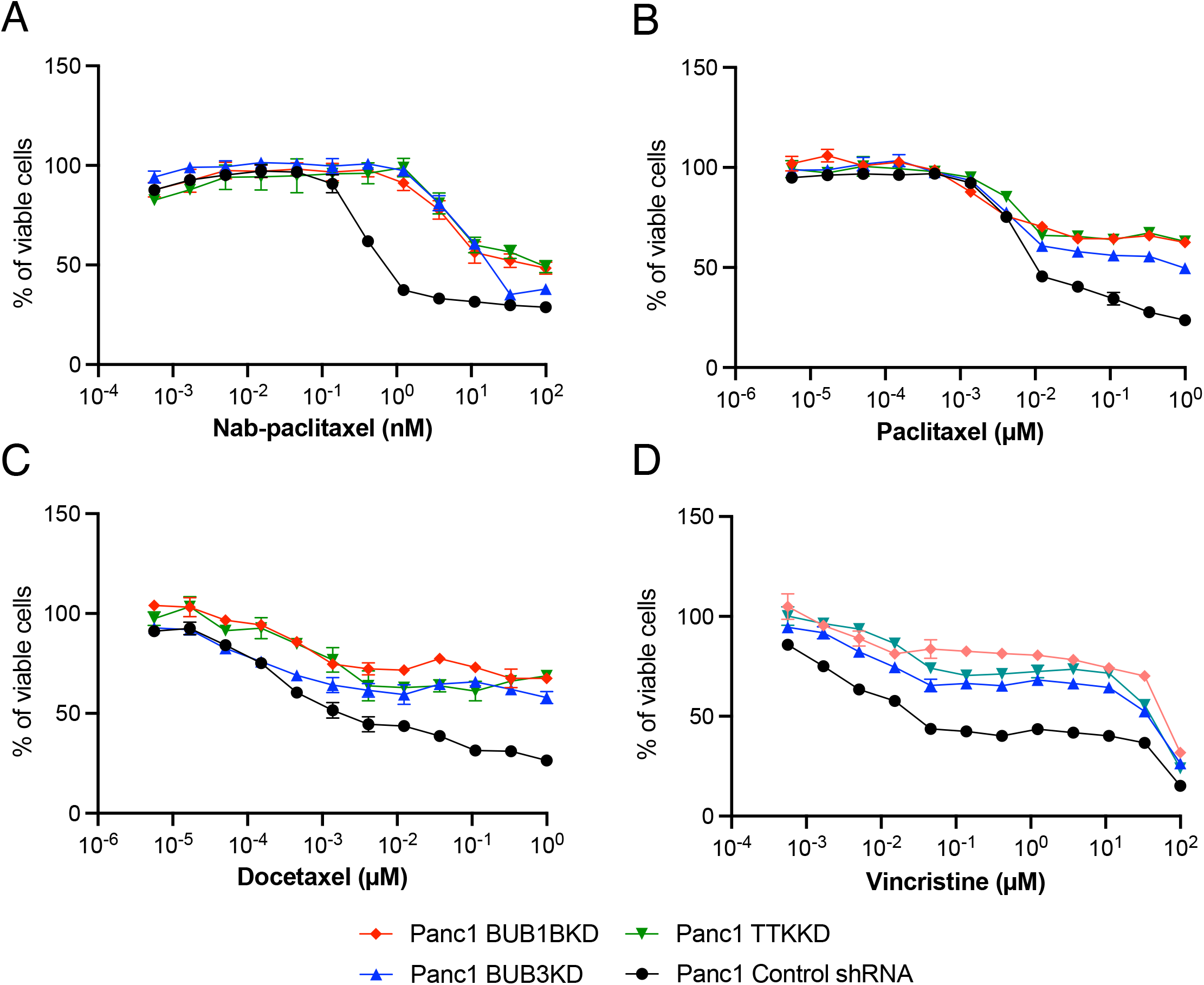
Reducing the activity of specific SAC genes diminishes the responsiveness of PANC-1 cells to Abraxene and other drugs that target microtubules. ShRNA-transfected PANC-1 cells were treated with (A) Nab-paclitaxel, (B) Paclitaxel, (C) Docetaxel, and (D) Vincristine for 96 hrs. Relative viable cell numbers were measured using the Cell Titer-Glo assay. Data represent three independent experiments with mean ± SEM.

### Inhibition of selected SAC genes decreases the cell proliferation of PANC-1 and creates aneuploidy

Impairment in the spindle assembly checkpoint (SAC) can lead to aneuploidy and chromosomal instability, increasing cancer risk^21^. Aneuploid cancer cells are less sensitive to multiple chemotherapeutic drugs^22,23^. Under normal culture conditions, PANC-1 BUB1B KD and TTK KD cells exhibit slower proliferation than control cells. However, in the presence of low-dose nab-paclitaxel (5 nM), BUB1B, TTK, or BUB3 knockdown cells exhibit faster cell proliferation than the NT control cells (Fig. 3A, B). We examined the distribution of cell cycle phases of shRNA-transfected PANC-1 cell lines in the presence or absence of nab-paclitaxel (1 μM, 24 hours) (Fig.3C). In the absence of nab-paclitaxel, PANC-1 cells with SAC gene knockdowns exhibited a major G2/M arrest compared to the control NT cells. In contrast, knockdown of BUB1B, TTK, or BUB3 cells showed little or no G2/M arrest when exposed to nab-paclitaxel compared to the control. However, drug treatment was associated with a significant increase in the tetraploid population in cells with SAC gene knockdown.

**Figure 3.**
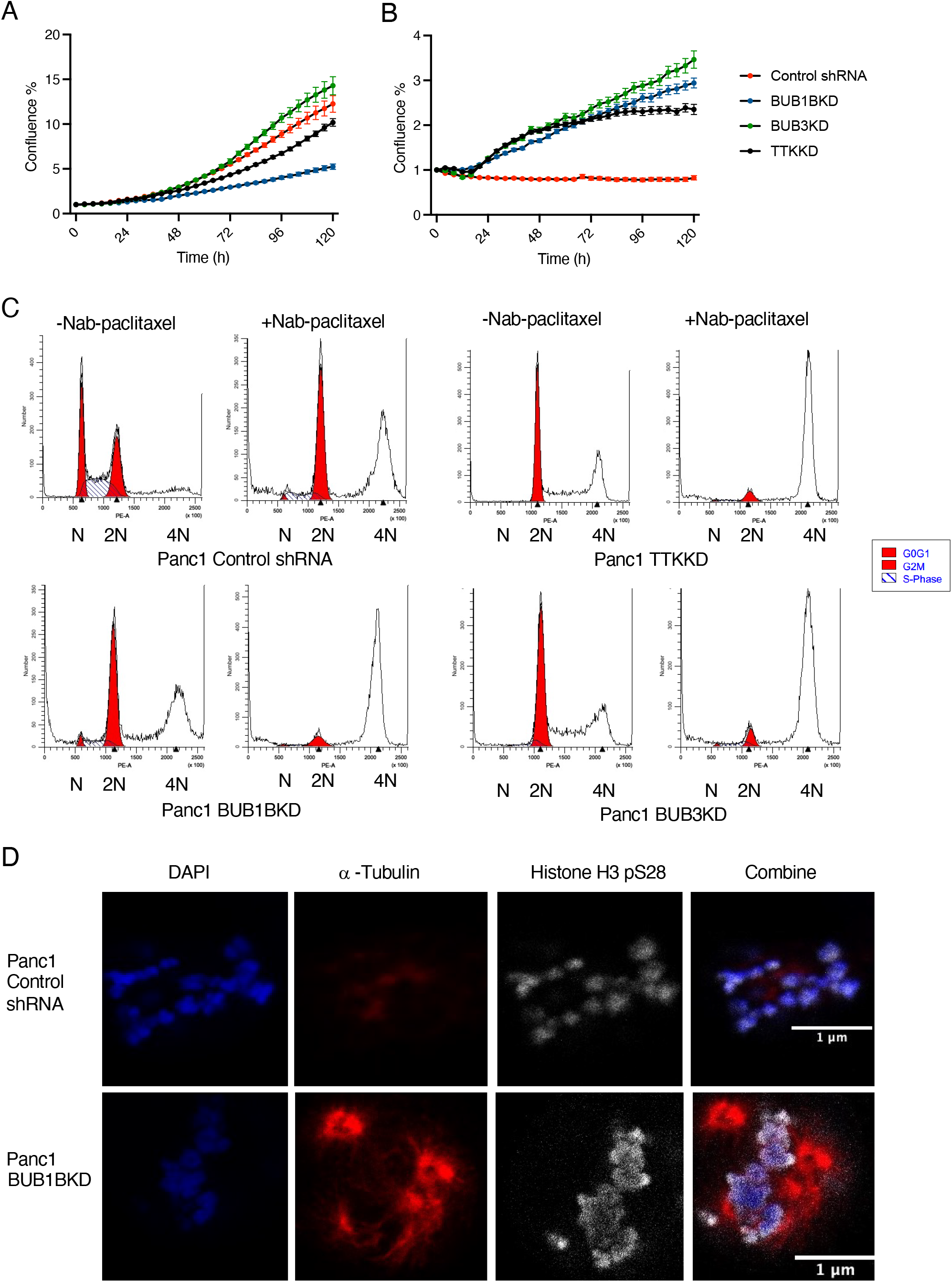
Blocking specific SAC genes leads to a decrease in PANC-1 cell proliferation and induces aneuploidy. Cell proliferation curves of shRNA-transfected PANC-1 cells cultured with (A) DMSO or (B) Nab-paclitaxel for 5 days. Data represent three biological replicates with mean ± SD. (C) Cell cycles of shRNA-transfected PANC-1 cells were measured with or without nab-paclitaxel for 24 hrs. The image represents one out of three biological replicates. (D) shRNA-transfected PANC-1 cells were treated with nab-paclitaxel and then immunostained for DAP1 (blue), Histone H3p28 (White), and α-tubulin (red). Scale bars: 1 μM.

To elucidate the impact of SAC gene expression deficiency on chromosome structure, we conducted a detailed examination of chromosome alignment during metaphase using immunofluorescence techniques (Fig. S3). To achieve this, cells were synchronized in metaphase and subjected to staining for α-tubulin, ds-DNA, and histone H3 pSer28. Notably, the knockdown of BUB1B and TTK resulted in abnormal chromosomal structures and misalignment during metaphase. Our analysis revealed a distinct alteration in chromosome size for cells with knockdowns of BUB1B, TTK, and BUB3, indicating a significant departure from the chromosomal characteristics observed in normal PANC-1 cells. The enlarged chromosomes observed in these knockdown cells suggest a potential link between the absence of crucial SAC genes and altered chromosomal morphology, possibly resulting from the failure of cells to divide properly. It has been reported that the cytotoxicity of paclitaxel is due to chromosome mis-segregation on multipolar spindles^24^. Utilizing time-lapse microscopy, we observed that paclitaxel indeed led to chromosomal misalignment and the formation of multipolar spindles (Fig. 3D). Following a 12-hr treatment with 1 nM nab-paclitaxel, both PANC-1 and PANC-1 BUB1BKD cells exhibited nuclear segregation. However, PANC-1 BUB1BKD cells displayed a notably higher incidence of multipolar spindles, accompanied by a more condensed signal in the presence of nab-paclitaxel. This suggests that the compromised SAC gene, particularly BUB1B, may overcome the effects of paclitaxel, further contributing to chromosomal stability and aberrant spindle formation.

### Clinical correlation of low SAC proteins with poor outcome in ovarian cancer

A Kaplan-Meier survival analysis was carried out to determine the relationship between the expression of BUB1B, TTK, and BUB3 and the overall survival and recurrence-free survival of ovarian cancer patients treated with paclitaxel. The findings showed that high expression levels of BUB1B, BUB3, or TTK were associated with better survival rates in ovarian cancer patients treated with paclitaxel compared to patients with low expression levels (Fig. S4).

### TTK inhibitors antagonize the cytotoxic effect of nab-paclitaxel in PANC-1

TTK is a SAC protein that plays a crucial role in cell proliferation and division. It is essential for proper chromosome alignment at the centromere during mitosis and centrosome duplication^25^. Many studies have shown a connection between high TTK expression and malignant progression in different types of cancer, including pancreas cancer, gastric cancer, colon cancer, clear cell renal cell carcinoma, prostate cancer, breast cancer, non-small-cell lung cancer, and medulloblastoma^26-33^.

Inhibition of TTK resulted in cells with mis-segregated chromosomes due to premature exit of mitosis^34^. Earlier studies showed that a few TTK inhibitors, such as MPI-0479605, suppressed the growth of cancer xenografts in immune-deficient mice. In contrast, others like MPS-BAY2b synergized with anti-mitotic drugs such as paclitaxel or vincristine to inhibit the growth of cancer xenografts in immune-deficient mice^35,36^. Several TTK inhibitors have entered clinical trials for advanced cancer patients. Two examples are BAY 1161909 (ClinicalTrials.gov identifier: NCT02138812) and BAY 1217389 (ClinicalTrials.gov identifier: NCT02366949). Other clinical trials have investigated whether inhibiting TTK can improve the efficacy of paclitaxel in treating solid tumors (see NCT03411161, NCT03328494, NCT02366949).

To investigate the impact of SAC protein kinase inhibitors in the presence of nab-paclitaxel, we chose two well-studied TTK inhibitors, BAY 1217389 and MPI-0479605. We found that these TTK inhibitors could counteract the cytotoxic effect of nab-paclitaxel (Fig. 4A, B). The multidomain protein kinases (BUB1) play a central role in the mitotic checkpoint for the spindle assembly, similar to BUB1B^37^. We next explored the impact of BAY 1816032. This specific inhibitor operates by selectively targeting BUB1 kinase, a key component of the SAC. Notably, the addition of BUB1 kinase inhibitor resulted in an increase in cell cytotoxicity with nab-paclitaxel, suggesting a potential synergistic effect with nab-paclitaxel (Fig. 4C). Unlike the findings from our CRISPR screens, this result suggests that targeting of SAC components is not universally associated with nab-paclitaxel resistance. Instead, selective inhibitors show the ability to counteract the cytotoxic effects induced by nab-paclitaxel in PANC-1 cells.

**Figure 4.**
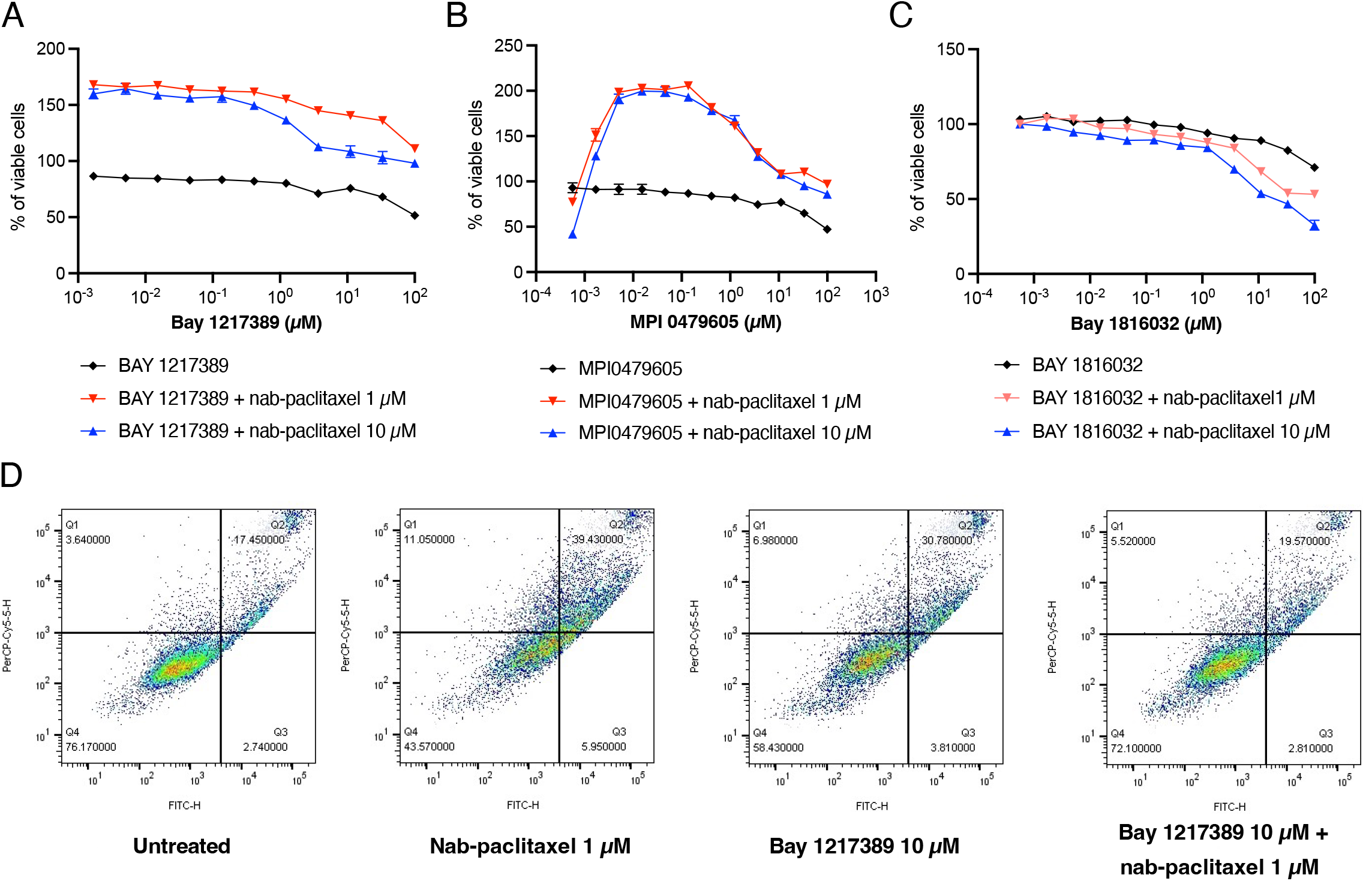
TTK inhibitors counteract the cytotoxic impact of nab-paclitaxel in PANC-1 cells. PANC-1 cells were treated with (A) BAY 1217389, (B) MPI0479605, and (C) BAY 1816032 for 72 hrs in the presence of a fixed concentration of nab-paclitaxel. Relative viable cell numbers were measured using the CellTiter-Glo assay. Data show three independent experiments with mean ± SEM. (D) Apoptotic death of PANC-1 cells was measured with BAY 1217389 or nab-paclitaxel for 48 hrs. Each image represents one out of three biological replicates.

We next examined whether the TTK inhibitors could potentially protect against nab-paclitaxel-induced apoptosis. PANC-1 cells were subjected to various conditions for 48 hrs, including 1 μM nab-paclitaxel alone, 10 μM BAY 1217389 alone, and the combination of 1 μM nab-paclitaxel and 10 μM BAY 1217389. The apoptotic response was assessed by flow cytometry quantification of Annexin V and propidium iodide (PI) staining. Treatment with either a single agent, nab-paclitaxel or BAY 1217389, demonstrated the capacity to induce apoptosis in PANC-1 cells. Strikingly, the combination of nab-paclitaxel and BAY 1217389 resulted in a significant reduction in the number of apoptotic cells compared to the individual treatments (Fig. 4D). The number of apoptotic cells after the combination treatment was similar to what was observed with untreated cells. Taken together, these results suggest that the TTK inhibitor may rescue the cells from the apoptotic effects of paclitaxel.

### BUB1B, BUB3, or TTK overexpression has no effect on the sensitivity of PANC-1 cells to nab-paclitaxel treatment

Overexpression of the SAC gene has been observed in various cancers, including pancreatic, thyroid, prostate, and liver cancers^38-41^. Our study found that overexpression of the three SAC genes (BUB1B, TTK, and BUB3) does not significantly affect sensitivity to nab-abraxane. We conducted cytotoxicity assays using Abraxane on PANC-1 cells that either overexpressed one of the three genes or had empty vectors. As shown in Fig 5A, there was no change in cell viability observed in PANC-1 cells that overexpressed BUB1B, TTK, or BUB3 compared to the control group. We examined the distribution of cell cycle phases of PCDNA-transfected PANC-1 cell lines in the presence or absence of nab-paclitaxel (Fig. 5B). Cells that overexpressed BUB1B, TTK, BUB3, or empty vectors were treated with 1 μM nab-paclitaxel for 24 hr. The PANC-1 cells that overexpressed SAC exhibited a longer G2/M phase compared to the control group. After the treatment with nab-paclitaxel, the SAC-overexpressing cells showed a significant increase in arrest during the S and G2/M phase, but few tetraploid cells were detected. The overexpression of BUB1B, TTK, and BUB3 was confirmed by measuring protein expression levels (Fig. S5).

**Figure 5.**
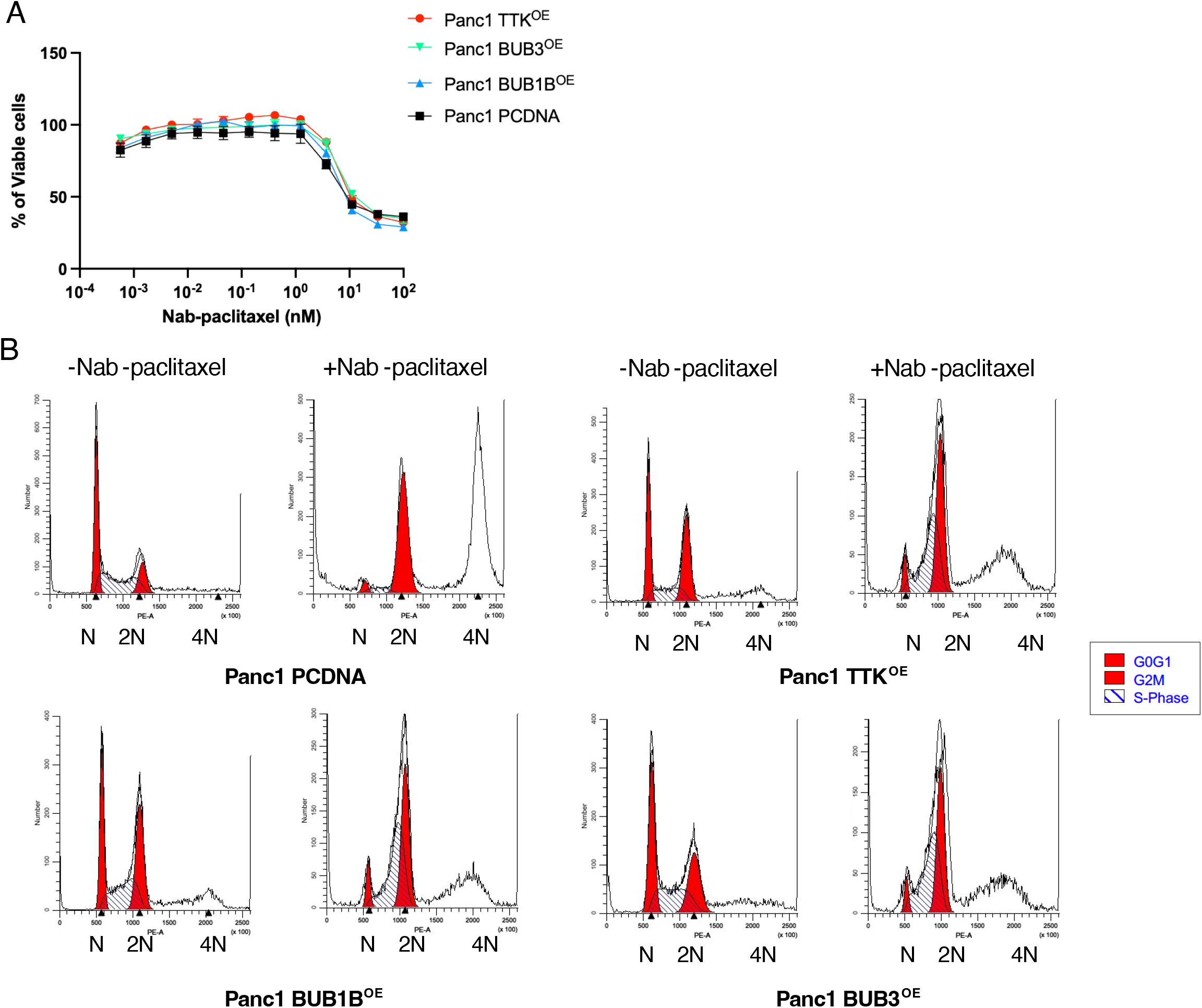
Overexpressing BUB1B, BUB3, or TTK does not impact the sensitivity of PANC-1 cells to nab-paclitaxel treatment. (A) pcDNA-transfected PANC-1 cells were treated with nab-paclitaxel for 96 hrs. Relative viable cell numbers were measured using the Cell Titer-Glo assay. Data shown represent three independent experiments with mean ±. (B) Cell cycles of PCDNA-transfected PANC-1 cells were measured with or without Abraxane for 24 hrs. The image represents one out of three biological replicates.

## Discussion

Paclitaxel remains a cornerstone in cancer treatment despite the advent of immune therapy and targeted small molecule inhibitors, yet its precise mechanism of action is still poorly understood. Our study delves into the intricacies of nab-paclitaxel resistance in a model of pancreatic adenocarcinoma, PANC-1, and pinpoints the pivotal role played by three SAC proteins—BUB1B, TTK, and BUB3—in determining sensitivity of a pancreatic cancer cell line to paclitaxel. Our results strongly suggest that the primary mechanism of action of paclitaxel in these cells is to prevent microtubules from attaching to kinetochores, resulting in activation of the spindle assembly checkpoint (SAC) and the failure of cells to divide. Our investigation reveals that disrupting the SAC, either through biological or chemical means, induces an increase in aneuploidy. This adaptive response facilitates the evasion of nab-paclitaxel-induced mitotic arrest. Conventionally, paclitaxel was thought to synergize with SAC inhibitors, particularly TTK. However, our results challenge this paradigm by demonstrating that TTK inhibitors impede the efficacy of nab-paclitaxel. This finding may help to explain the setbacks observed in several phase 1 clinical trials, such as with the SAC protein kinase inhibitors NCT03411161, NCT03328494, and NCT02366949. The evolving landscape of personalized medicine in oncology emphasizes tailoring treatments based on the genetic variability of tumors. Our study contributes useful data to support this approach, concluding that patients exhibiting elevated expression of the BUB1B, TTK, and BUB3 genes are likely to derive greater benefit from nab-paclitaxel/paclitaxel regimens. This nuanced understanding of genetic variations provides a foundation for more effective and personalized cancer treatment strategies.

During cell division processes such as mitosis and meiosis, the spindle assembly checkpoint (SAC) plays a crucial role in maintaining the stability of the genome. It does so by delaying cell division until accurate chromosome segregation can be guaranteed. BUB1B, BUB3, and TTK are key proteins involved in SAC signaling and stable attachment of kinetochores to spindle microtubules^42,43^. BUB1B/BUBR1 (BUB1 mitotic checkpoint serine/threonine kinase B) is a member of the SAC protein family^44^. BUB1B interacts directly with Cdc20, BUB3, and MAD2, constituting the mitotic checkpoint complex. This complex inhibits the activity of the anaphase-promoting complex or cyclostome (APC/C)^45^. The TTK (threonine tyrosine kinase) protein kinase, also known as monopolar spindle 1 (MPS1), is essential for SAC protein localization and mitotic checkpoint complex (MCC) production^43^. The SAC genes mentioned earlier are essential for cell growth and survival. We have confirmed that culturing stable BUB1B/BUB3/TTK CRISPR knockouts is impossible and in the studies reported here, we used stable shRNA knockdowns that retained small amounts of the SAC proteins to allow long-term cell viability.

Studies have shown that low levels of BUB1B and BUB3 are linked to poor prognosis of carcinomas^46-48^. Inhibiting the activity of BUB1B, TTK, and BUB3 can impede cancer cell proliferation, migration, and invasion^49-51^. Furthermore, reducing BUB1B and TTK levels or inhibiting their kinase activity causes a significant loss of chromosomes, ultimately leading to apoptotic cell death^52,53^. BUB1B overexpression drives cancer progression and recurrence, promoting anchorage-independent survival and facilitating lung adenocarcinoma metastasis^54^. In extrahepatic cholangiocarcinoma, BUB1B regulates proliferation and invasiveness^55^. Elevated TTK expression is associated with aggressive phenotypes in triple-negative breast cancer cells^56^. Abnormal Bub3 expression leads to spindle gate defects, chromosomal instability, and aneuploidy, crucial for malignancy genesis, progression, and metastasis^57^.

The paradox, of course, is that impeding an essential function such as the SAC allows cells to survive a cytotoxic treatment. These genetic results are confirmed with the studies using SAC protein kinase inhibitors, though not all inhibitors show this effect, either because they have off-target effects that are cytotoxic in the presence of paclitaxel, or because they are less potent in inhibiting the SAC. Our findings reinforce previous observations in the literature^58-60^ and indicate that sensitivity to drugs is a complex phenomenon that depends on the precise genetic and epigenetic environment. Whether PANC-1 cells will prove to be a suitable model for studying the drug sensitivity of pancreatic cancer remains to be determined.

Another surprising result of these studies is that the CRISPR-Cas9 screen we utilized did not find that expression of ABCB1 (P-glycoprotein) is a major mechanism of resistance to nab-paclitaxel, whereas it is known (and we have confirmed) that overexpression of P-gp can reduce sensitivity to nab-paclitaxel^61^. Furthermore, in other CRISPR-Cas9 screens we have conducted with other drugs that are substrates for efflux by P-glycoprotein, ABCB1 is the major mechanism by which cells become resistant (data not shown). It has been speculated that nab-paclitaxel, a complex with nanoparticle albumen, may be able to enter cells via receptor-mediated endocytosis^62^, which might, under some circumstances, bypass P-glycoprotein-mediated efflux of paclitaxel. Our results are consistent with the possibility that nab-paclitaxel is a less likely substrate for P-glycoprotein than paclitaxel alone, at least in PANC-1 cells.

There is still much to learn about how complex cancer cell biology influences the response of cancer cells to chemotherapy. Despite being one of our most effective anti-cancer drugs, paclitaxel remains something of an enigma, and we will not be able to optimize its use to treat cancer until we understand more completely how it kills cells and how cells circumvent its toxicity.

## Acknowledgements

We would like to express our gratitude to Dan Sackett, NICHD, for his scientific advice, as well as to Michael Kruhlak, at the NCI Microscopy Core for his technical assistance. We thank Rob Robey for valuable advice and helpful suggestions, and George Leiman for editorial assistance.

## Author contributions

G.A. designed the Crispr screen in consultation with M.M.G. P.M. designed all other experiments and acquired data. P.M. and A.V.M. analyzed data. P.M. wrote the manuscript. M.M.G. revised the manuscript, with the help of the other authors. All authors read and approved the final version of the paper.

## Funding

This research was funded by the Intramural Research Program of National Cancer Institute in the National Institutes of Health.

## Data Availability

All of the data generated through this research are contained within the manuscript and associated supplemental files.

## Competing interests

The authors declare no competing interests.

## Notes

### Competing Interest Statement

The authors have declared no competing interest.

