## Supplemental Figures S1-S5 for "A whole-genome CRISPR screen identifies the spindle accessory checkpoint as a locus of nab-paclitaxel resistance in pancreatic cancer cells"

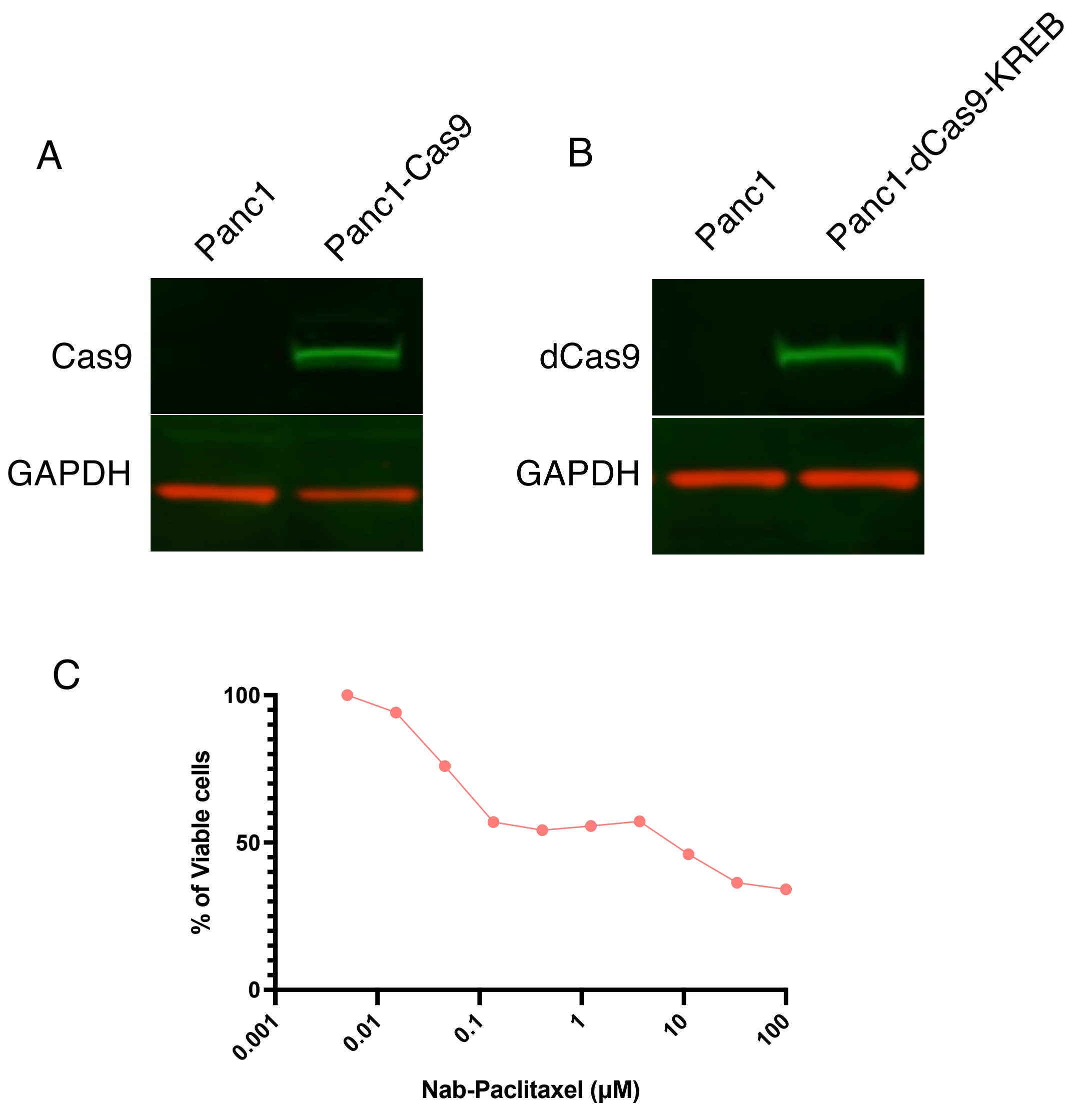


**Fig. S1.** The stable expression of Cas 9 (A) and dCas9-KREB (B) is confirmed by Western blot. GAPDH is used as a control. (C) The proliferation of Panc1 cells is significantly suppressed by 10 μM nab-paclitaxel treatment, ensuring an effective selection pressure for the genetic screen.

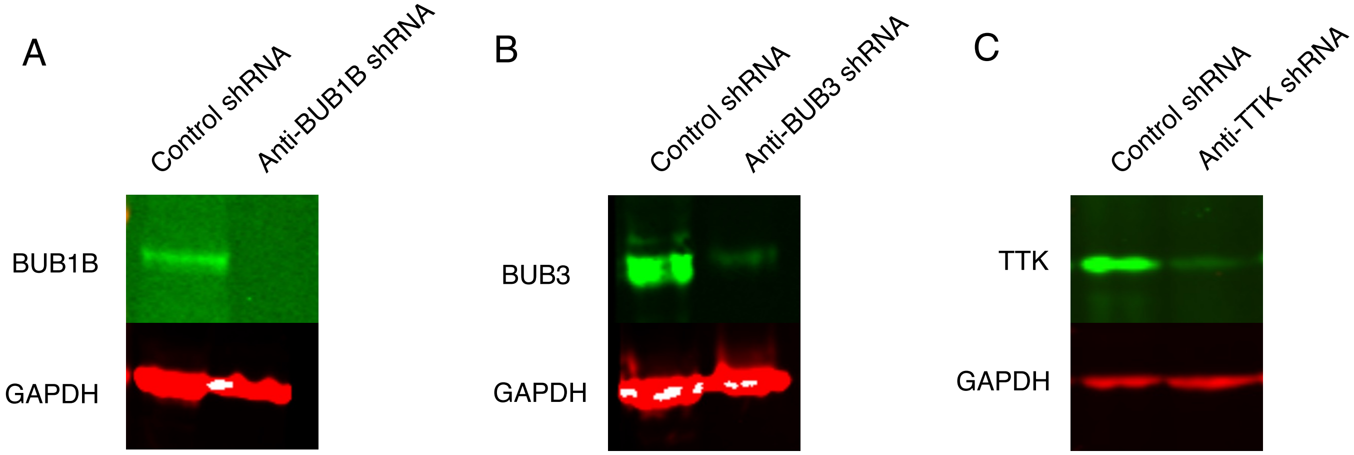


**Figure S2.** Knockdown of (A) BUB1B, (B) BUB3, and (C) TTK by lentiviral-based shRNA is confirmed at the protein level by Western blot. GAPDH was used as a loading control.

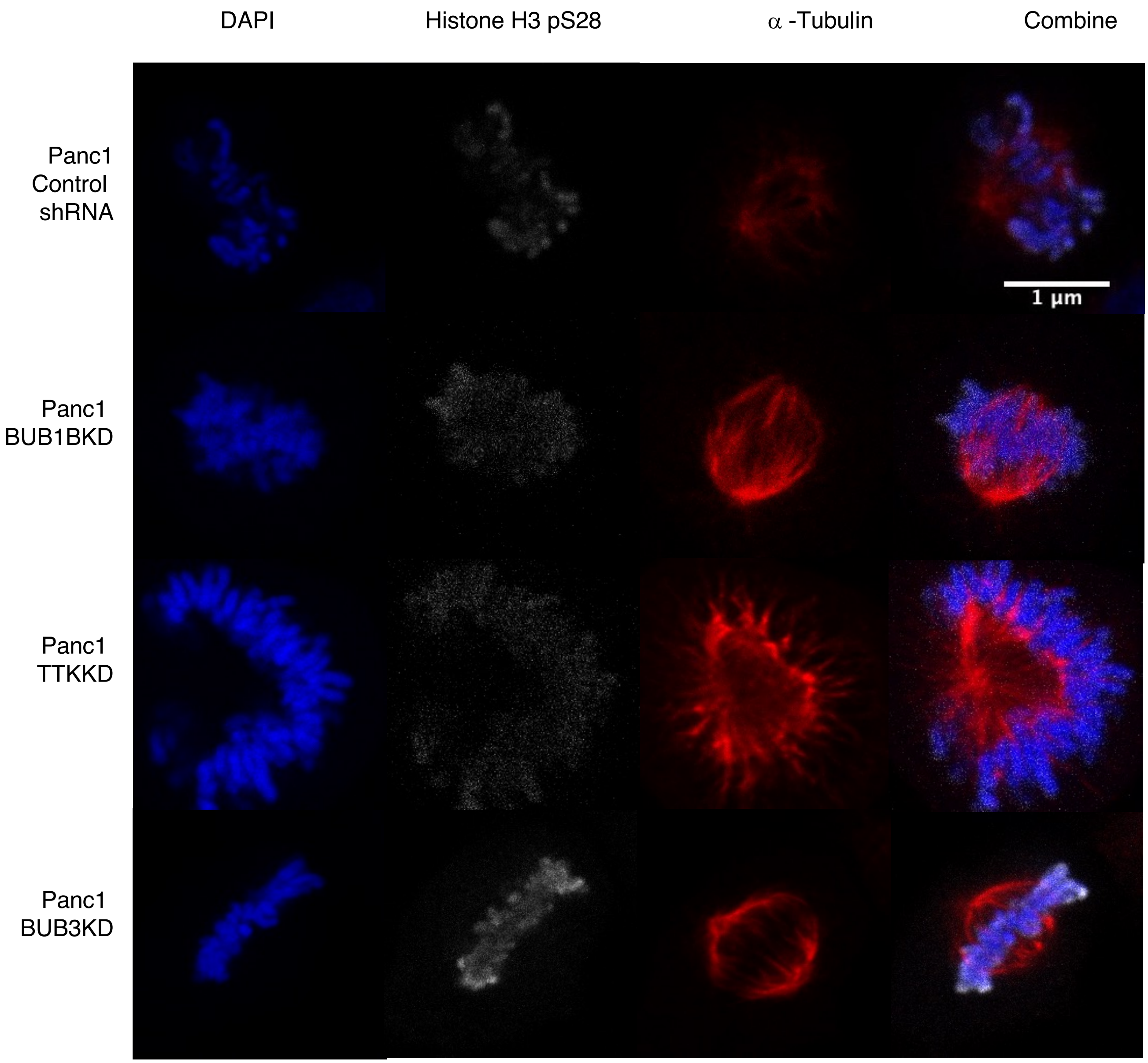


**Figure S3.** shRNA-transfected Panc1 cells were immunostained for DAP1 (blue), Histone H3p28 (white), and α-tubulin (red). Scale bars: 1 μM.

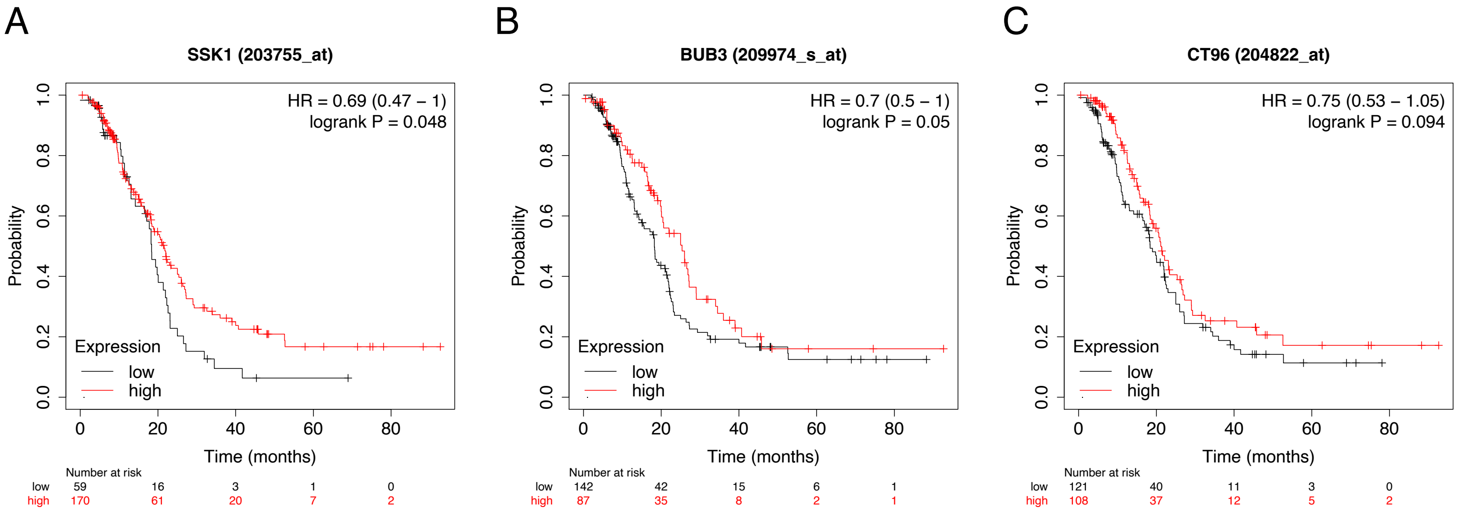


**Figure S4.** KM Plotter database showing the correlation between the expression of (A) BUB1B (SSK1), (B) BUB3, and (C) TTK (CT96) with overall survival. Survival curves were plotted based on the mRNA values from approximately 250 ovarian cancer patients treated with paclitaxel.

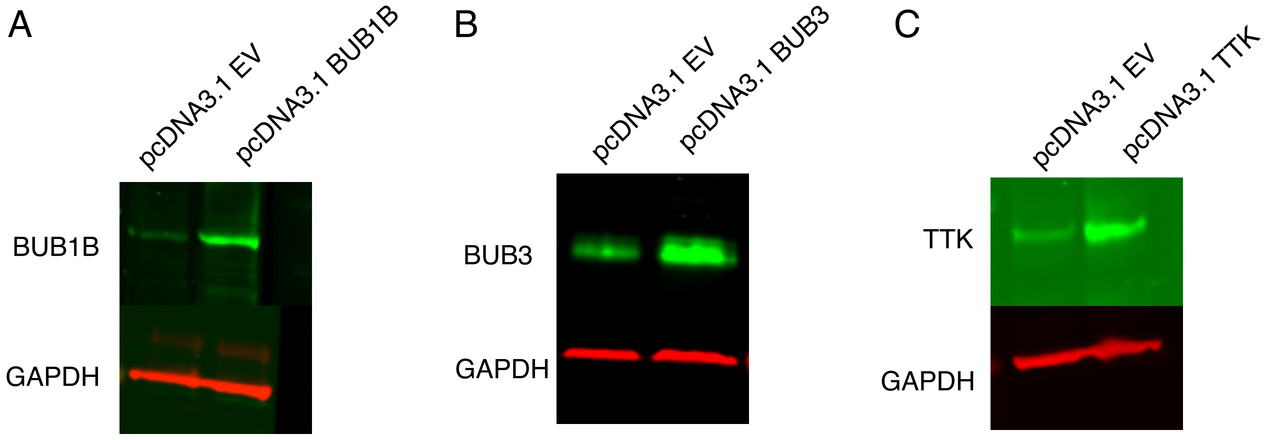


**Figure S5.** Overexpression of (A) BUB1B (B) BUB3 (C) TTK by PCDNA is confirmed at the protein level by Western blot. GAPDH is used as a loading control.
